# The neurophysiology of healthy and pathological aging: A comprehensive systematic review

**DOI:** 10.1101/2024.08.06.606817

**Authors:** Gemma Fernández-Rubio, Peter Vuust, Morten L. Kringelbach, Leonardo Bonetti

## Abstract

As the population of older adults grows, so does the prevalence of neurocognitive disorders such as mild cognitive impairment (MCI) and dementia. While biochemical, genetic, and neuroimaging biomarkers have accelerated early detection and diagnosis, neurophysiological measures are absent from daily medical use. Electroencephalography (EEG) and magnetoencephalography (MEG) are two non-invasive techniques that measure neurophysiological signals in the brain and convey information about signal strength at different frequency bands, event-related activity, signal complexity, and temporal correlation between spatially remote brain regions. Here we conducted a pre-registered, comprehensive systematic review of 942 studies using EEG, MEG, and combined MEG and EEG to characterise the neurophysiology of healthy aging, MCI, and dementia under resting-state and task conditions. To complement our search, we also reviewed 51 past reviews in the field. Relevant features of these papers were extracted to present a detailed overview of the current state of evidence. Overall, neurophysiological measures show great promise as diagnostic tools and could prove invaluable in predicting healthy and pathological aging trajectories. However, to reach this potential in clinical practice, it is crucial to adopt longitudinal designs, standardise methodologies, and identify biomarkers at the individual rather than group level.

## Introduction

Due to advances in healthcare and standards of living, the population of older adults is rapidly growing. In 2022, there were an estimated 771 million people over the age of 65 (10% of total population), and this figure is expected to soar to 1.6 billion (16%) by 2050 ^1^. In parallel, age is a major risk factor for neurocognitive disorders such as mild cognitive impairment (MCI) and dementia. This situation brings about pressing public and economic concerns, underscoring the need to develop effective treatments, along with screening and diagnostic tools that can predict healthy and pathological aging trajectories.

The natural aging process is associated with a gradual decline in memory, attention, and other cognitive functions while activities of daily living remain intact ^2^. However, if deficits become noticeable and persistent in one or more cognitive functions, individuals may be diagnosed with MCI, a transitional stage to dementia ^3^. In this case, cognitive symptoms significantly interfere with daily activities, imposing significant burden and distress on caregivers ^4^. MCI and dementia are multifaceted neurocognitive disorders that can be due to a number of aetiologies ^5^. The most common form is Alzheimer’s disease dementia (ADD), which contributes to approximately 60-70% of cases, followed by vascular dementia (VD), dementia with Lewy bodies (DLB), frontotemporal dementia (FTD), and mixed dementia ^6^. The global number of cases of dementia are projected to rise from 57 million in 2020 to 153 million in 2050, with AD-related dementias creating an estimated $16.9 trillion economic burden in 2050 ^7,8^. While no known cure exists, current treatments can temporarily slow down the progression of symptoms, especially when administered at initial stages of the disease. For this reason, early diagnosis is crucial to delay the onset of dementia and lower healthcare costs^9^.

Biochemical, genetic, and neuroimaging biomarkers have made great strides toward early detection. For example, novel cerebrospinal fluid (CSF) biomarkers may differentiate ADD from FTD and DLB ^10^. In the case of genetic biomarkers, a dementia gene panel supplemented by testing for the *C9orf72* expansion is the most suitable method for detecting most dementia forms ^11^. Structural and molecular neuroimaging techniques, such as magnetic resonance imaging (MRI) and positron emission tomography (PET), are widely used in clinical practice^12^. For example, biomarkers of ADD include amyloid-beta and tau pathology, structural changes in the hippocampus, and cerebrovascular flow ^13^. On the other hand, neurophysiological measures, which provide high-dimensional multivariate brain data, have not been introduced in daily medical practice. Moreover, studies that correlate these measures with current biochemical, genetic, and neuroimaging biomarkers are scarce ^14,15^.

Neurophysiological signals are primarily recorded using electroencephalography (EEG) and magnetoencephalography (MEG), two non-invasive techniques that can characterise changes in brain activity based on the electrochemical activity produced by thousands of neurons ^16^. EEG can detect tangential and radial components from current sources near the skull, while MEG is sensitive to magnetic fields originating from tangential components only. Magnetic fields are less distorted than electrical signals by brain tissue, conferring higher spatial resolution to MEG than EEG systems and allowing for more accurate source reconstruction, particularly when combined with individual structural brain images from MRI^17^. Metrics of EEG and MEG can provide information about the strength of the signal at different frequency bands (spectral analysis), the changes in activity related to specific events (event-related analysis), the complexity of brain signals (complexity analysis), and the temporal correlation between spatially remote signals (functional connectivity analysis) ^16^.

Neurophysiological techniques have captured significant attention from the scientific community, leading to a broad consensus regarding their effectiveness to detect pathophysiological mechanisms of cognitive decline in both healthy and pathological aging ^18–21^. While previous reviews provided valuable insights, they primarily focused on the potential of neurophysiology to differentiate ADD patients from healthy controls, often neglecting the context of healthy aging. Additionally, these reviews often concentrated on specific technical aspects of neurophysiological signals or narrow subtopics within this extensive field. As a result, they typically lack breadth, encompassing a range of 50 - 100 studies each ^14,16,22–27^.

The overarching goal of this study is therefore to provide a comprehensive, large-scale systematic review that synthesises evidence from the complete spectrum of EEG, MEG, and combined EEG and MEG (M/EEG) literature on both healthy and pathological aging, including all forms of dementia and MCI. After screening 6156 unique articles, we extracted data from 942 studies and sought to (1) characterise demographic trends in publication, (2) present the classical features of these studies in terms of experimental setup, population, and data analysis, (3) summarise key results, and (4) examine limitations and provide recommendations for the field. By integrating a vast array of studies, our review provides a thorough inspection of the up-to-date body of evidence on the neurophysiology of healthy and pathological aging.

## Methods

### Search strategy, study identification, and pre-registration

We performed a systematic review of publications reporting neurophysiological measures (EEG, MEG, and M/EEG) in healthy aging, mild cognitive impairment (MCI), and dementia using PubMed, Scopus, and PsychInfo. The search (14 December 2023) included MeSH terms (“electroencephalography”, “magnetoencephalography”, “healthy aging”, “Alzheimer disease”, “dementia”) and key words (“mild cognitive impairment”, “early stages of Alzheimer disease”, “early stages of dementia”, “early onset dementia”, “early onset Alzheimer disease”, “aging”, “resting-state”, “task”, “task complexity”, “predictive progression”, “predictive progression of dementia”, “prediction”, “functional connectivity”, “structural connectivity”). Two reviewers (GFR and LB) independently screened by title and abstract on Covidence ^28^ and selected articles for full-text review and data extraction. Discrepancies were resolved through discussion with MLK.

We prospectively registered a protocol for our systematic review in the International Prospective Register of Systematic Reviews (PROSPERO) on 28 November 2022 with registration number CRD42022375928.

### Eligibility criteria

To be eligible for inclusion, studies had to meet the following criteria: (1) peer-reviewed publications in English, (2) on healthy aging, MCI, or dementia, and (3) using EEG, MEG, or M/EEG. No year or location restrictions were imposed. The following exclusion criteria were applied: (1) sleep studies, (2) treatment studies, (3) studies on epileptiform activity, and (4) review studies. Review studies were included in a separate, additional review.

### Data extraction and analysis

From the studies that met our inclusion criteria, we extracted the following features: (1) first author, (2) last author, (3) title, (4) year of publication, (5) journal, (6) Digital Object Identifier (DOI) or PubMed identifier (PMID), (7) affiliation of first author, (8) country of affiliation, (9) source of funding, (10) study design, (11) study duration (only applicable to longitudinal studies), (12) neurophysiology technique, (13) additional measure(s), (14) paradigm I, (15) paradigm II, (16) population, (17) sample size, (18) number of females, (19) age, (20) analysis, (21) feature analysed, (22) machine learning, (23) source reconstruction, and (24) key results. This information is reported in **Supplementary Table 1** (ordered alphabetically by first author). Descriptive statistics and plots from selected features were computed in RStudio ^29^, while panels were created using Adobe Illustrator ^30^.

To complement the results from this systematic review, we extracted the following information from past reviews on the topic: (1) first author, (2) last author, (3) title, (4) year of publication, (5) journal, (6) DOI or PMID, (7) affiliation of first author, (8) country of affiliation, (9) type of review, (10) number of reviewed articles, (11) time period of reviewed articles, (12) neurophysiology technique, (13) additional measure(s), (14) paradigm, (15) population, and (16) key results. This information is reported in **Supplementary Table 2** (ordered alphabetically order by first author). Statistical analyses were not performed, as features from previous reviews were extracted solely for the purpose of complementing our systematic review.

## Results and Discussion

The literature search produced 8195 articles from PubMed (*n* = 4090), Scopus (*n* = 2056), and PsychInfo (*n* = 2049). After duplicates were removed, 6156 unique articles were screened for title and abstract. A total of 1238 articles met the screening criteria and were assessed for eligibility. After removing further duplicates (*n* = 9), unavailable publications (*n* = 41), previous reviews (*n* = 51), studies that did not meet the eligibility criteria (*n* = 90), and conference proceedings (*n* = 105), we reviewed 942 articles. **Figure 1** depicts the literature search and selection process, while information on included studies is reported in **Supplementary Table 1**. To complement our results, we performed a minor review of past reviews on the topic, as reported in **Supplementary Table 2**.

**Figure 1.**
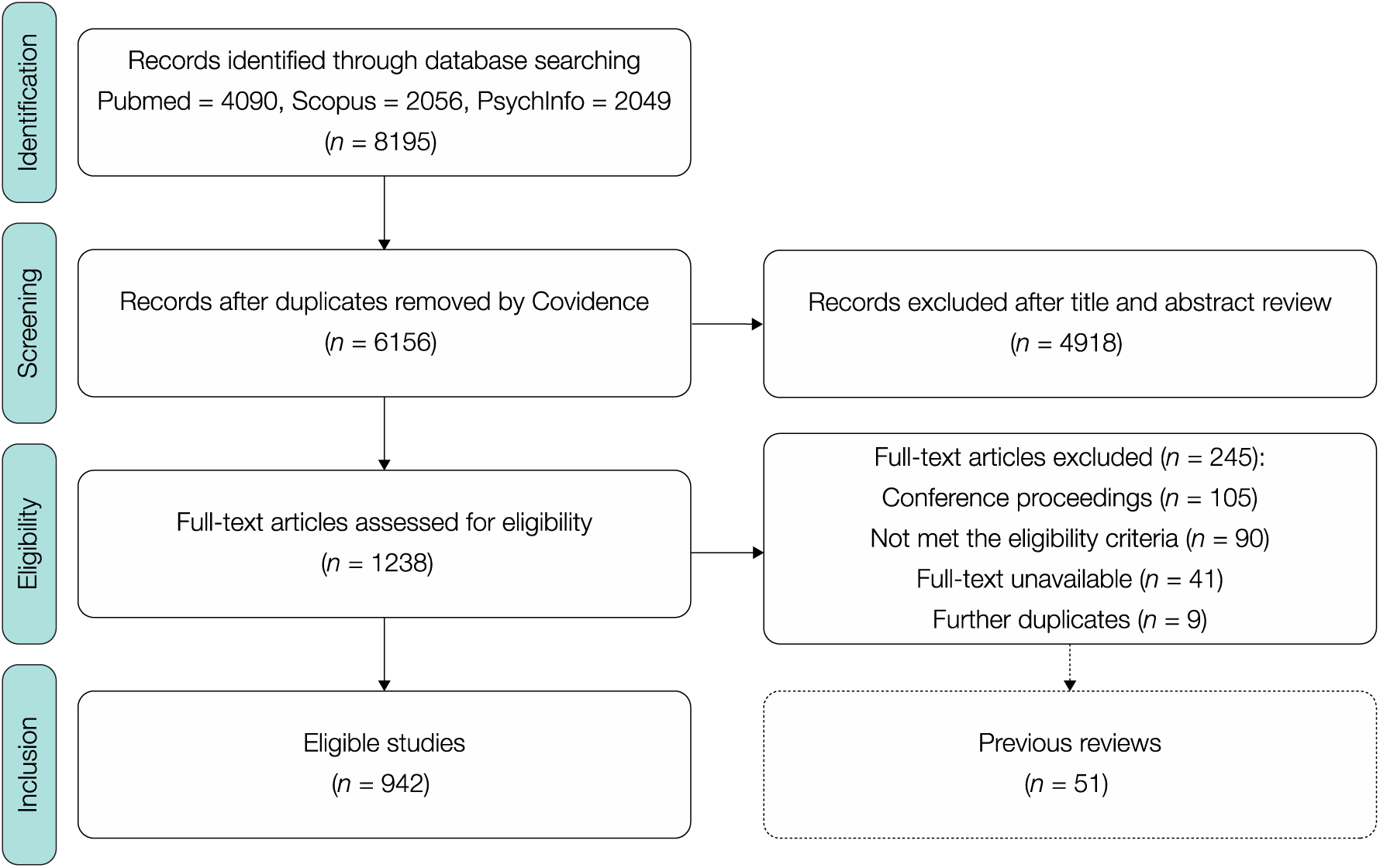
Selection process and results from literature search. After record identification, screening, and eligibility assessment, we included 942 studies in our systematic review and reported their main features in **Supplementary Table 1**. As illustrated by the dotted box, an additional systematic review was conducted on previous reviews and outlined in **Supplementary Table 2**.

### Demographics

We extracted the date and country of publication of 942 articles to determine the demographic characteristics of studies on the neurophysiology of healthy aging, MCI, and dementia.

Regarding the temporal distribution, there is a clear and consistent increase in the number of publications per year, particularly in the last decade. The most dramatic increase was experienced between 2021 and 2023, with publications rising from 54 in 2021 (5.73% of total reviewed articles) to 136 in 2022 (14.44%) and 138 in 2023 (14.65%). While this recent surge is consistent with pattern changes in scientific publishing associated with the effects of the COVID-19 pandemic ^31^, the general rising trend reflects a swift progress in the field of neurophysiology and a growing interest in understanding the neurophysiological mechanisms of healthy and pathological aging. The temporal distribution of reviewed articles is presented in **Figure 2a**, while the publication year of each article is displayed on column E (“Year”) of **Supplementary Table 1**.

**Figure 2.**
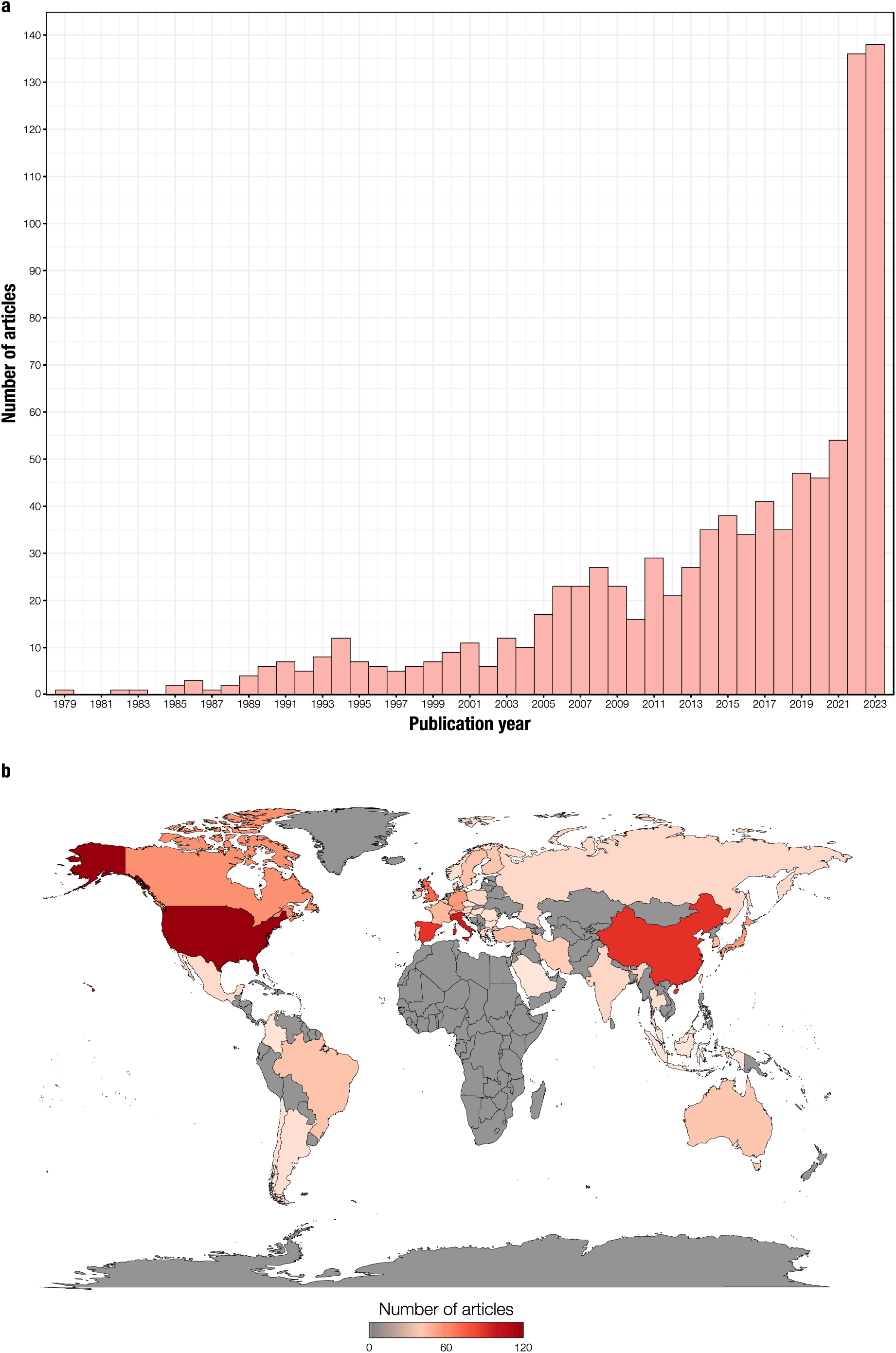
Demographics of reviewed articles. **a** – Number of reviewed articles per publication year (n = 942). **b** – Number of reviewed articles per country (n = 942). The geographic distribution was determined based on the affiliation of the first author.

We determined the geographic distribution of reviewed articles based on the affiliation of the first author. The top 10 publishing countries were USA (12.95% of total reviewed articles), Italy (10.72%), China (9.34%), Spain (9.13%), The Netherlands (6.58%), UK (6.47%), Canada (4.46%), Germany (4.03%), Japan (3.93%), and Switzerland (2.65%). Seven of these countries are among the top 10 publishing countries from 1996 to 2023, according to the SCImago Journal Rank ^32^. As **Figure 2b** illustrates, there is an overrepresentation of high-income countries, followed by publications from middle-income countries, and no studies from low-income countries. This in stark contrast with the most recent report from the World Health Organization (WHO) on the distribution of dementia worldwide, which states that over 60% of affected individuals live in middle- and low-income countries ^6^. On **Supplementary Table 1**, columns H (“Affiliation”) and I (“Country”) provide information on each reviewed article.

### Experimental setup

To estimate the experimental characteristics of the 942 articles reviewed, we extracted three key features: (1) study design, (2) neurophysiology technique, and (3) paradigm.

The majority of articles reviewed employed a cross-sectional design (*n* = 870) and this trend was stable over the years (see **Figures 3a** and **3b**). The ratio of cross-sectional to longitudinal studies was consistently 75% or higher, except in 1985 (one cross-sectional and one longitudinal article, 50%) and in 1987 (one longitudinal study, 100%), as depicted in **Figure 3c**. A handful of studies reported the lack of longitudinal data as a limitation ^33,34^ and most authors pointed to the benefits of a longitudinal design and provided suggestions for future studies. This is particularly relevant for monitoring conversion from preclinical to prodromal to dementia stages and for improving the diagnostic utility of neurophysiological measures ^25^. However, a longitudinal design is more costly and effortful than a cross-sectional one, which could explain the lack of longitudinal studies reviewed here. The study design of each reviewed article can be found in Column K (“Design”) of **Supplementary Table 1**.

**Figure 3.**
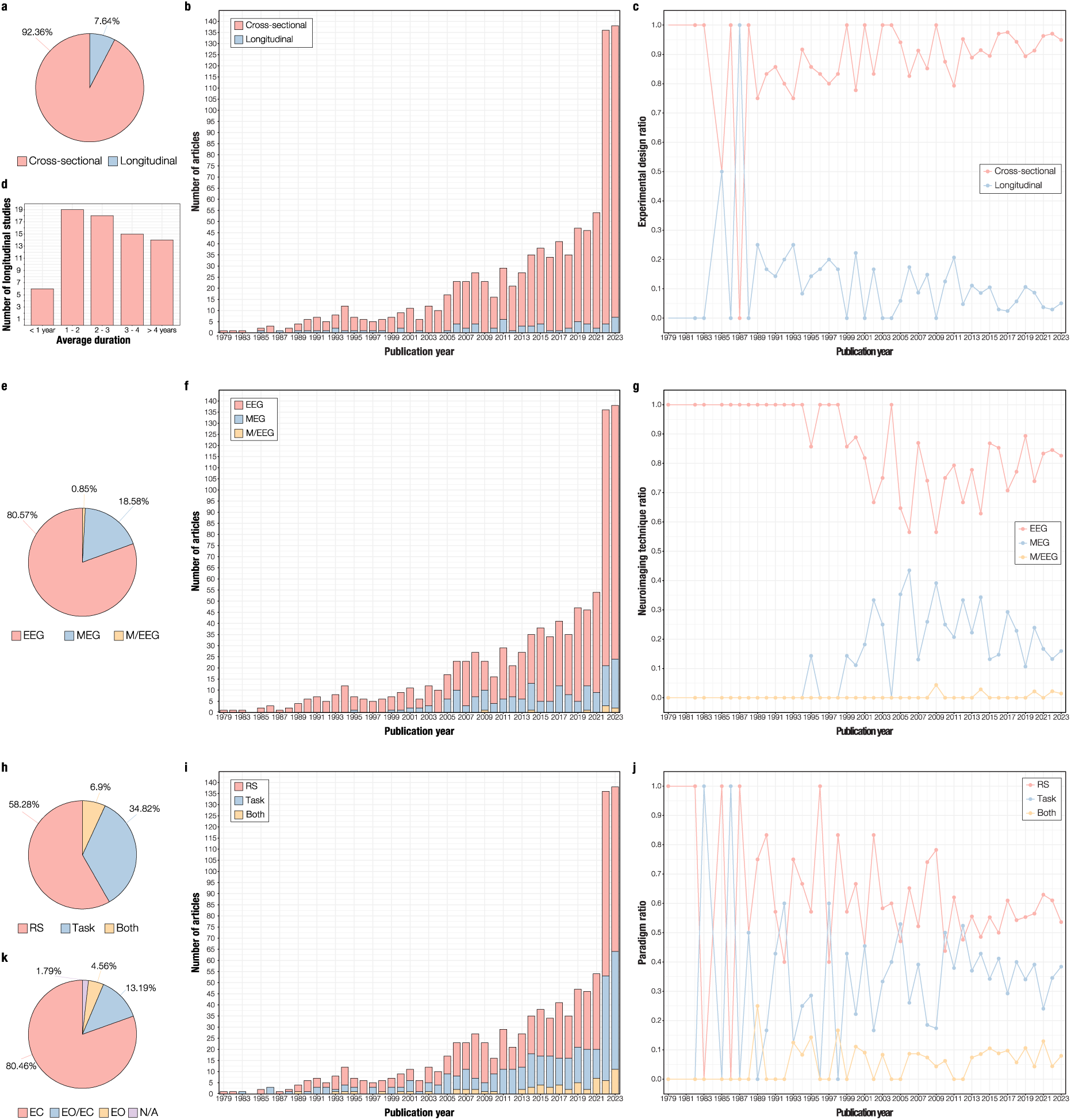
Experimental setup of reviewed articles. **a** – Percentage of cross-sectional and longitudinal articles reviewed (n = 942). **b** – Number of cross-sectional and longitudinal articles reviewed per publication year. **c** – Ratio of cross-sectional to longitudinal articles reviewed per publication year. **d** – Duration of longitudinal studies (n = 72), classified into one of five lengths: less than 1 year, between 1 and 2 years, between 2 and 3 years, between 3 and 4 years, and over 4 years. **e** – Percentage of electroencephalography (EEG), magnetoencephalography (MEG), and combined EEG and MEG (M/EEG) articles reviewed (n = 942). **f** – Number of EEG, MEG, and M/EEG articles reviewed per publication year. **g** – Ratio of EEG, MEG, and M/EEG articles reviewed per publication year. **h** – Percentage of articles reviewed measuring brain activity during resting-state (RS), task, or both (n = 942). **i** – Number of RS, task, and combined RS and task articles reviewed per publication year. **j** – Ratio of RS, task, and combined RS and task articles reviewed per publication year. k – Percentage of studies measuring RS during eyes closed (EC), alternating eyes open and eyes closed (EO/EC), eyes open (EO), and not specified (N/A).

We found that 51.39% of longitudinal studies (*n* = 72) reported an average duration of 1-2 and 2 - 3 years, which are standard and feasible periods of time (see **Figure 3d**). However, such limited time windows could restrict tracking the progression of dementia symptoms, which can take up to 5 years to develop in cognitively active older adults ^35^. The longest study reviewed here collected resting-state EEG data from patients with young onset ADD and young onset FTD for a median of 10 years (3 - 20 years range), and focused on differences between the two groups in slow wave patterns and epileptiform activity ^36^. The duration of longitudinal studies is outlined in Column L (“Duration”) of **Supplementary Table 1**.

Regarding neurophysiological measures, 80.57% of articles reviewed employed EEG (see **Figure 3e**). This trend was stable until the late 1990s, when the number of studies employing MEG systems started rising steadily (**Figure 3f**). However, the proportion of EEG to MEG and M/EEG studies was always greater than 50% over publication years (**Figure 3g**). Regarding M/EEG systems, only 8 studies (0.85%) reported using this technique. The prevalent use of EEG can be linked to its longer history and lower cost than MEG systems, which are less available worldwide. Results from MEG and EEG studies tend to be coherent, particularly regarding spectral features such as slowing patterns in ADD patients ^24^. However, metrics of functional connectivity and network architecture in the AD continuum can be inconsistent between the two modalities ^15^. Furthermore, while MEG excels in discriminating MCI patients from healthy controls, EEG is superior in differentiating ADD patients from controls ^26^. A lack of homogeneity in sample characteristics, source localization methods, and data analysis could explain the discrepancies, and highlights the urgency of a shared methodological approach between EEG and MEG groups. Information on each reviewed article is reported in Column M (“Neurophysiology technique”) of **Supplementary Table 1**.

Additional measures included neuropsychological testing (*n* = 156), MRI (*n* = 80), apolipoprotein E genotype (*n* = 16), CSF biomarkers (*n* = 14), and PET (*n* = 13). Overall, these measures were used to characterise experimental groups and complement neurophysiological techniques. Details for each reviewed article are reported in Column N (“Additional measure(s)”) of **Supplementary Table 1**.

With respect to experimental paradigms, over half of the reviewed articles report measuring EEG, MEG, or M/EEG during resting-state, followed by task-based studies, and studies employing both resting-state and tasks (see **Figure 3h**). Studies measuring resting-state exhibit a steady and constant increase over publication years, task-based studies present the sharpest rise among the three experimental paradigms, particularly from 2007, and studies employing both resting-state and tasks are consistently fewer (see **Figures 3i** and **3j**). Experimental paradigms are reported in columns O (“Paradigm I”) and P (“Paradigm II”) of **Supplementary Table 1**.

The most widespread experimental condition was resting awake with eyes closed, which represents a standardised and low-cost procedure that can be readily implemented in neurophysiological recordings of older adults ^22^ (see **Figure 3k**). One potential limitation of this approach is the increase of alpha band activity in the spectral power ^23^. Furthermore, resting-state paradigms provide information on background activity only and do not probe brain activity during specific states that are relevant to the study of healthy and pathological aging (e.g., memory or attentional processes).

We identified 234 different experimental tasks in 393 studies, with 19.17% of them measuring automatic processing of auditory and visual change detection using oddball paradigms. Importantly, in 2022, an Expert Panel from the Electrophysiology Professional Interest Area of the Alzheimer’s Association concluded that event-related oscillations (EROs) in EEG are altered in ADD, Parkinson’s disease dementia (PDD), DLB, and cerebrovascular diseases, and recommended the use of oddball paradigms to compare event-related theta and delta responses in patients with MCI, ADD, PDD, DLB, and vascular cognitive impairment ^14^. This represents a major development in the establishment of neurophysiological biomarkers of dementia. Tasks measuring working memory, inhibitory control, and short-term memory were also among the 10 most commonly used, providing insights into the mechanisms underlying affected cognitive functions in older adults. The 10 most common tasks are presented in **Table 1**, while a complete report of all tasks and their frequency is presented in **Supplementary Table 3**.

**Table 1.** Ten most common experimental tasks.

| Task | Modality | Measure | No. articles reviewed (%) |
| --- | --- | --- | --- |
| Oddball paradigm | Auditory | Auditory ERP/Fs | 60 (13.54) |
| Oddball paradigm | Visual | Visual ERP/Fs | 25 (5.63) |
| N-back task | Visual | Working memory | 18 (4.04) |
| Go/no-go task | Visual | Inhibitory control | 13 (2.93) |
| Sternberg letter-probe task | Visual | Working memory | 13 (2.93) |
| Delayed match-to-sample task | Visual | Working memory | 12 (2.70) |
| Passive stimulation | Auditory | Auditory ERP/Fs | 11 (2.47) |
| Mental arithmetic task | N/A | Working memory | 9 (2.03) |
| Short-term memory task | Visual | Short-term memory | 5 (1.13) |
| Working memory task | Visual | Working memory | 5 (1.13) |

**Note.** ERP/Fs = Event-related potentials/fields

### Population

To describe the population of the reviewed studies, we focused on three broad categories (healthy aging, MCI, and dementia) and characterised the frequency and subgroups within each category.

The majority of studies reviewed comprised at least one group with dementia (see **Figure 4a**). This trend was stable until 2000, when the number of studies on the neurophysiology of healthy aging and MCI started growing until a certain balance between the three population groups was achieved in 2023, with 42.03% of studies on dementia, 29.71% on healthy aging, and 28.26% on MCI (see **Figures 4b** and **4c**).

**Figure 4.**
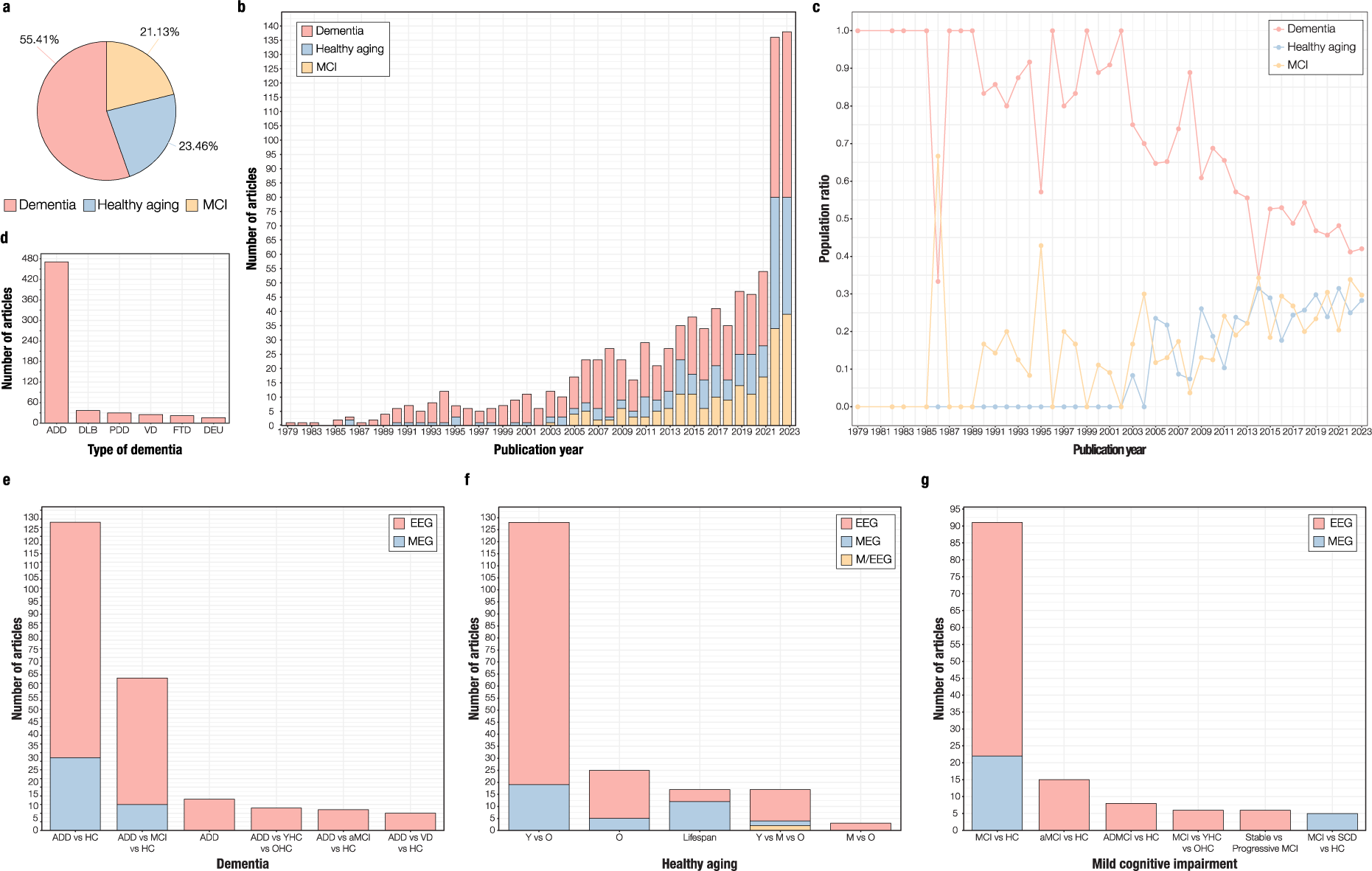
Population characteristics of reviewed articles. **a** – Percentage of dementia, healthy aging, and mild cognitive impairment (MCI) articles reviewed (n = 942). Population was classified as (1) dementia (i.e., at least one experimental group comprises dementia patients), (2) MCI (i.e., at least one experimental group comprises MCI patients), or (3) healthy aging (i.e., none of the experimental groups comprises dementia nor MCI patients). **b** – Number of dementia, healthy aging, and MCI articles reviewed per publication year. **c** – Ratio of dementia, healthy aging, and MCI articles reviewed per publication year. **d** – Number of studies including at least one group with Alzheimer’s disease dementia (ADD), dementia with Lewy bodies (DLB), Parkinson’s disease dementia (PDD), vascular dementia (VD), frontotemporal dementia (FTD), and dementia of aetiology unknown (DEU). **e** – Top 5 dementia groups per neurophysiology technique. The x-axis displays the most prevalent sample designs: Alzheimer’s disease dementia (ADD) patients versus healthy controls (“ADD vs HC”), ADD patients versus MCI patients versus healthy controls (“ADD vs MCI vs HC”), ADD patients only (“ADD”), ADD patients versus younger healthy controls versus older healthy controls (“ADD vs YHC vs OHC”), ADD patients versus amnestic MCI patients versus healthy controls (“ADD vs aMCI vs HC”), and ADD patients versus vascular dementia patients versus healthy controls (“ADD vs VD vs HC”). The y-axis presents the number of articles per group and neurophysiology technique: electroencephalography (EEG), magnetoencephalography (MEG), and combined EEG and MEG (M/EEG). **f** – Top 5 healthy aging groups per neurophysiology technique. The x-axis shows the most prevalent healthy aging sample designs: younger versus older adults (“Y vs O”), older adults only (“O”), across lifespan (“Lifespan”), younger versus middle-aged vs older adults (“Y vs M vs O”), and middle-aged versus older adults (“M vs O”). The y-axis displays the number of articles per group and neurophysiology technique. **g** – Top 5 MCI groups per neurophysiology technique. The x-axis depicts the most prevalent MCI sample designs: MCI patients versus healthy controls (“MCI vs HC”), amnestic MCI patients versus healthy controls (“aMCI vs HC”), patients with MCI due to Alzheimer’s disease versus healthy controls (“ADMCI vs HC”), MCI patients versus younger healthy controls versus older healthy controls (“MCI vs YHC vs OHC”), stable versus progressive MCI patients (“Stable vs Progressive MCI”), and MCI patients versus subjective cognitive decline individuals versus healthy controls (“MCI vs SCD vs HC”). The y-axis presents the number of articles per group and neurophysiology technique.

Widespread interest in the neurophysiology of dementia is expected, given its high incidence, economic burden, and predicted growth ^7,8^. However, in the last two decades, attention was diverted to the prodromal stages of dementia, with a sharp rise in the number of studies on MCI and on healthy aging. This not only illustrates increasing trends in life expectancy, but a collective objective from the neurophysiological field to portray pre-clinical features and predict conversion to dementia.

Among dementia studies, the majority recruited at least one group with ADD, followed by studies including patients with DLB, PDD, VD, FTD, and dementia of unknown aetiology (see **Figure 4d**). This is akin to the distribution of prevalence of different dementia forms ^6^. Correspondingly, the most common sample designs compared ADD patients with healthy controls, and ADD patients with MCI patients and healthy controls. Interestingly, a handful of studies recruited ADD patients only (see **Figure 4e**) or healthy older adults only (see **Figure 4f**). Healthy aging studies largely contrasted groups of young and older adults, and MCI studies primarily investigated differences between MCI patients and healthy controls (see **Figure 4g**). A full description of each population group, sample size, number of females, and age is beyond the scope of this study. However, information on these features can be found in **Supplementary Table 1** on columns Q (“Population”), R (“N”), S (“F”), and T (“Age”), respectively.

### Data analysis

We extracted information on the primary data analysis methods employed by each study and classified them into five categories, mirroring previous focalised reviews ^15,23^. The categories were (1) spectral analysis, (2) event-related analysis, (3) functional connectivity analysis, (4) complexity analysis, and (5) other (i.e., analytical methods that did not fit into one of the other categories). Additionally, we collected basic information on the use of source reconstruction and machine learning methods.

Reviewed studies extracted and analysed mainly spectral, event-related, and functional connectivity features, followed by mixed (i.e., two or more features from different categories), complexity, and other features (see **Figure 5a**). This trend was stable over publication years, particularly from 2000 onwards (see **Figures 5b** and **5c**). Information on methods of data analysis for each article is provided in columns U (“Analysis”) and V (“Features”) of **Supplementary Table 1**. To ease readability, we grouped similar features together and assigned broad terms to describe them.

**Figure 5.**
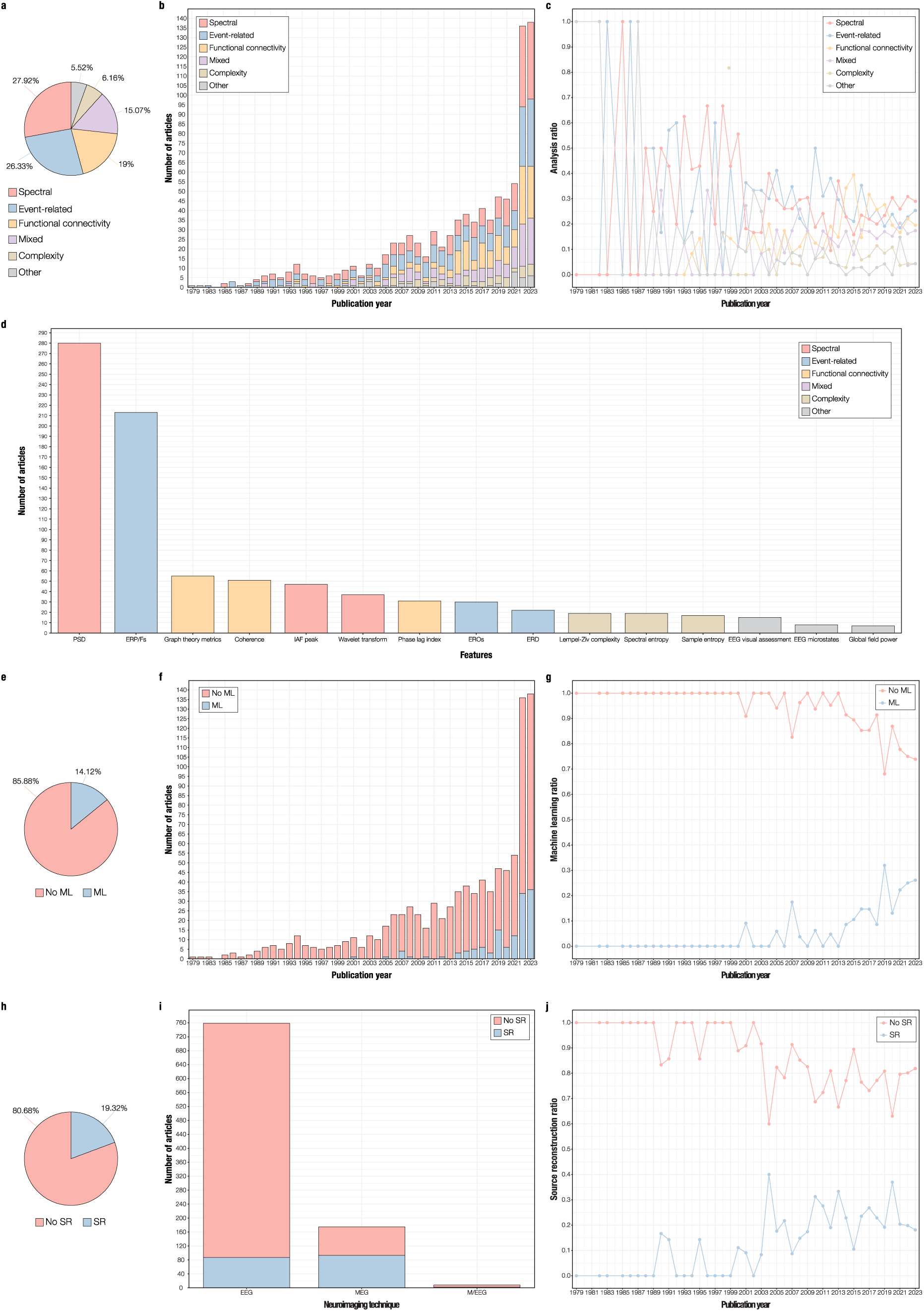
Analytical methods of reviewed articles. **a** – Percentage of reviewed studies performing analysis of spectral, event-related, functional connectivity, complexity, or other features (n = 942). The term “mixed” refers to the use of two or more of the aforementioned features. **b** – Number of articles reviewed performing analysis of spectral, event-related, functional connectivity, complexity, mixed, or other features per publication year. **c** – Ratio of reviewed articles performing analysis of spectral, event-related, functional connectivity, complexity, mixed, or other features per publication year. **d** – Top 3 features per category of analysis. The x-axis displays the three most used features for each type of analysis (spectral, event-related, functional connectivity, complexity, and other). The y-axis presents the total number of articles reviewed per feature. PSD = Power spectral density; ERP/Fs = even-related potentials/fields; IAF peak = individual alpha frequency peak; EROs = event-related oscillations; ERD = event-related desynchronization; EEG = electroencephalography. **e** – Percentage of reviewed articles using machine learning (ML) techniques (n = 942). **f** – Number of articles reviewed using ML techniques per publication year. **g** – Ratio of articles reviewed using ML techniques per publication year. **h** – Percentage of reviewed articles using source reconstruction (SR) techniques (n = 942). **i** – Number of articles reviewed using SR per neurophysiology technique: electroencephalography (EEG), magnetoencephalography (MEG), and combined EEG and MEG (M/EEG). **j** – Ratio of articles reviewed using SR techniques per publication year.

The leading neurophysiological metrics (*n* = 493) were power spectral density (PSD) and event-related potentials/fields (ERP/Fs), which is coherent with our results regarding the distribution of resting-state and task-based studies (see **Figure 5d**). PSD-based features describe the linear distribution of the spectral power over different frequency bands and have been proposed to be used as baseline for novel analytical methods ^23^. ERP/Fs, on the other hand, refer to neurophysiological signals that are time-locked to specific events such as sensory stimulation or participant responses. We illustrate the top three features per category of analysis in **Figure 5d** and compile all features and their frequency of occurrence per analysis category in **Supplementary Table 4**.

Among spectral features, individual alpha frequency (IAF) peak and wavelet parameters were extensively computed. The IAF peak estimates the maximum power density peak at the extended alpha frequency band, while the term “wavelet transform” was used to describe time-frequency analyses of EEG and MEG signals ^23^. For analysis of event-related features, EROs, which quantify the power dynamics of oscillations coupled to specific events, and event-related desynchronization (ERD), a measure of the amplitude of frequency-related ongoing oscillatory responses, were also extensively computed, in line with recommendations from Güntekin et al. (2022) ^14^. Regarding functional connectivity analysis, studies primarily calculated graph theory metrics, coherence, and phase lag index. Graph theory metrics are not a measure of functional connectivity per se, but were included as a separate feature within the functional connectivity category due to their prominence in the field, as previously denoted by Miraglia et al. (2017) ^37^. The top three complexity features were Lempel-Ziv complexity, spectral entropy, and sample entropy. Regarding other features, a number of studies visually inspected EEG signals, analysed EEG microstates, or computed global field power (GFP), a measure of generalised EEG amplitude.

A minority of studies employed machine learning from 2001 onwards and particularly in the last 10 years (see **Figures 5e, 5f**, and **5g**). Overall, these studies aimed to classify population groups by employing multiple neurophysiological markers embedded in machine learning algorithms. A recent review by Rossini et al. (2022) ^38^ showcased the value of combining machine learning with graph theory techniques for the early diagnosis of AD using EEG. Due to the considerable diversity of machine learning methods found in this review, we suggest performing a specialised systematic review to track down the most effective machine learning algorithms and features. The use of machine learning techniques is reported for each article in column W (“Machine learning”) of **Supplementary Table 1**.

Finally, a minority of studies applied source reconstruction methods (see **Figure 5h**). The use of source reconstruction techniques increased in the last two decades and was primarily linked to MEG studies (53.13%) compared to EEG (11.46%) and M/EEG studies (25%), as depicted in **Figures 5i** and **5j**. Analogous to machine learning methods, we observed a wide array of source reconstruction techniques which are beyond the scope of the current review. The use of source reconstruction methods is outlined in column X (“Source reconstruction”) of **Supplementary Table 1**.

### Key results

We provide an essential and qualitative summary of the main results from the 942 articles reviewed. Specifically, we focused on findings that were most consistent among studies, bypassing individual results that did not contribute to a comprehensive and integrative picture. The summary was arranged into five main categories following the analysis labels introduced in the previous section. Individual results from each study are reported in column Y (“Key results”) of **Supplementary Table 1**, while results from previous reviews are outlined in **Supplementary Table 2**.

1. *Spectral markers of healthy and pathological aging.* In line with previous reviews, we found that MCI and dementia diagnoses are associated with signs of power spectrum “slowing” ^39–42^. Typically, slowing of EEG and MEG signals involves a decrease of absolute and relative power in high frequencies (alpha and beta, 8 - 30 Hz) and a power increase in low frequencies (delta and theta, 0.5 - 8 Hz) ^27,43^. Power spectrum slowing has been observed in ADD ^40,44^, DLB ^45^, PDD ^46^, and MCI patients ^42^. Reductions of the IAF peak are also reported in ADD and MCI patients compared to healthy controls ^47,48^. Similarly, in healthy aging, older adults present lower alpha power and IAF peak than younger individuals ^49–52^.
2. *Event-related markers of healthy and pathological aging.* In terms of ERP/F components, the reviewed studies showed smaller N400 effects ^53–57^ and smaller P300 amplitude in healthy older versus younger adults ^58–60^, as measured in a variety of active memory and inhibitory control tasks. Decreased mismatch negativity effects indexing automatic memory processes were also observed in ADD, VD, and MCI patients compared to controls ^61–65^, and in healthy older versus younger individuals ^58,66^. ERP/F components were recorded during auditory or visual oddball paradigms, facilitating result generalization. In fact, the only two meta-analyses found among previous reviews concluded that ADD and MCI patients present longer N200 latencies than healthy controls ^67^ and that ADD patients exhibit longer P300 latencies than MCI patients and controls ^68^. With respect to EROs and ERD, results substantially differed between studies, presumably due to the diversity of tasks used. However, we did observe an age-related increase in movement-related beta desynchronization and beta bursts during sensorimotor tasks ^69–71^.
3. *Functional connectivity markers of healthy and pathological aging.* One of the most consistent findings among functional connectivity markers was related to coherence, a measure that captures the phase consistency between spatially remote signals, such as two brain regions or two M/EEG channels. Reviewed studies showed reductions in coherence in patients with ADD ^72–76^, DLB ^77^, PDD ^78,79^, and MCI ^75,80^, as well as in older healthy adults ^81^. Such reduction in coherence was observed in different frequency bands and between a vast array of brain regions and M/EEG channels for different studies. Arguably, future research should highlight the key brain regions and frequency bands that mostly contribute to these differences in coherence. We found that MCI is associated with reduced phase lag index in the alpha band across different brain regions and M/EEG channels ^82–84^. Additionally, phase locking value is altered in healthy aging ^85–87^. However, results are not particularly consistent regarding the aging effect as well as the spectral and spatial distribution of these changes. Regarding graph theory metrics, we observed reduced characteristic path length in ADD patients, especially in the lower alpha band ^88^, decreased clustering coefficient in ADD patients in occipital regions ^89^ and VD patients in parietal regions ^90^, and increased small world index in the theta band in MCI patients ^91^. For a detailed report on EEG graph theory markers of healthy and pathological aging, we recommend the review by Miraglia et al. (2017) ^37^.
4. *Complexity markers of healthy and pathological aging.* Overall, reviewed studies highlighted a reduction of EEG and MEG signal complexity in ADD and MCI patients compared to healthy controls. A variety of metrics computed across the entire brain data were employed, particularly Lempel-Ziv complexity ^92–94^. Studies also found decreased entropy in several pathologies, such as ADD ^95^, VD ^96^, and MCI ^97^, which was also reported across the whole brain data. Taken together, these measures reveal a pattern of reduced complexity in the neurophysiology of pathological aging. Interestingly, this trend did not emerge in healthy aging, as greater entropy values were observed in older than in younger individuals, underlying the potential of entropy metrics to discern healthy from pathological aging ^98,99^.
5. *Other neurophysiological markers of healthy and pathological aging.* Beyond these key four categories of results, several individual findings were recorded which still share certain commonalities. For example, in line with key results of spectral markers of aging, studies performing visual assessments of EEG signals covering the entire scalp found that dementia patients present signs of EEG slowing compared to healthy controls ^100,101^. Additionally, several studies reported broad differences in coverage rate and occurrence frequency of a variety of EEG microstates between healthy and pathological populations ^102–104^ and between MCI and dementia patients ^105,106^. However, such differences were not always consistent between studies and call for future research. Finally, in terms of GFP, studies showed that increased whole-brain GFP in delta and theta frequency bands and reduced GFP in the alpha band is characteristic of ADD patients ^107–109^.

### Limitations and recommendations for future research

With global increases in longevity, neurophysiological measures provide a valuable tool to probe the brain mechanisms of healthy and pathological aging. However, upon inspection of the substantial body of literature, we encountered significant caveats and intrinsic limitations:

1. Together with the growing interest in the neurophysiology of healthy and pathological aging, there is an overrepresentation of publications from high-income countries. This is a long-standing challenge in the scientific community with considerable impact on the validation of biomarkers of dementia, as 60% of individuals with dementia live in middle- and low-income countries ^6^.
2. While cross-sectional studies have made outstanding contributions to the development of novel neurophysiological metrics, the dearth of longitudinal studies is alarming (7.64% of reviewed studies). Furthermore, the average length of reviewed studies was largely under 3 years, rendering short time windows for the prediction and monitoring of the development of neurocognitive disorders in healthy older adults.
3. Due to their lower cost and widespread availability, EEG systems have been more extensively used than MEG systems. This hinders our ability to conduct source reconstruction and raises the issue of compatibility between the two techniques, as they measure related but different components of electrochemical activity in the brain ^17^. Differences in sample characteristics and methodologies can further impact coherence of results, diminishing their prospects as diagnostic tools of dementia.
4. Recordings during resting with eyes closed in quiet vigilance have become the standard practice in EEG and MEG studies of healthy and pathological aging due to their easy implementation, followed by task-based conditions. However, the consistency of results between these very different experimental conditions has been largely unexplored, as evidenced by the lack of studies employing both conditions (6.9% of reviewed studies). Additionally, compared to resting awake with eyes open, eyes-closed conditions are associated with an increase in alpha power, which could affect the interpretation of results on power levels and brain dynamics at rest ^110^.
5. The emphasis on the neurophysiology of ADD is inevitably linked to its higher prevalence in the dementia population ^6^ but also leaves a substantial gap in the literature on other forms of dementia and MCI.
6. While the search of novel neurophysiological markers is promising, an overwhelming amount of EEG and MEG metrics were noted during data extraction, hampering a summary of the results and an effective comparison between different metrics. Overall, it appears that many novel metrics have been generated, leading to a disparity of results that do not provide a comprehensive theory of the neurophysiology of aging. As a matter of fact, among 51 past reviews, only two were meta-analyses. While these provided relevant results regarding differences in the latencies of specific auditory ERP components (N200 and P300) between ADD, MCI, and healthy individuals ^67,68^, the heterogeneity of metrics found here did not allow for direct quantitative comparisons of other findings.
7. Interestingly, although the reviewed articles often claimed to seek neurophysiological biomerkers for clinical purposes, their primary goal was to validate discrimination tools at the group level, as previously observed in a focalised systematic review by Cassani et al. (2018) ^23^. This shifts the focus away from screening and prediction at the individual level, which is instead necessary for developing screening tools that can be used in practical, real-world scenarios.

Based on the reviewed literature, we also provide a series of recommendations:

1. We strongly recommend longitudinal designs incorporating multiple neurophysiology recordings using EEG, MEG, or M/EEG. These measures should be complemented by established metrics of biochemical, genetic, neuroimaging, and neuropsychological data. This approach aims to assess and screen healthy older adults, validating candidate neurophysiological biomarkers and their link to established measures that could predict the onset of dementia. Given the gradual progression of the disease, we propose extending the average duration of these studies to a minimum of 5 years, with a preference for extending up to 10 years ^35^. We also emphasise that such transformative change should involve not only researchers but the entire scientific community, including foundations, stakeholders, and investors. Currently, most research grants, especially those for young researchers, provide funding for a relatively short duration (e.g., 5 years), which is why it is crucial to award different types of grants. For instance, the same amount of funding could be spread over 10 years instead of 5, or a process for extending existing grants for promising longitudinal designs could be implemented. While such funding mechanisms are not entirely absent, they require significant improvement. Similarly, the publication sector should more actively reward and encourage the development of longitudinal designs, potentially publishing interim findings to show the progression of the project in real time. This approach would increase transparency and, collectively, provide more useful and practical information suitable for real-life, clinical applications.
2. While several studies have generated large datasets using resting state conditions (particularly with eyes closed) and oddball paradigms, much remains to be done in task-based research. Although some excellent studies exist, future research should further explore tasks that mimic real-life scenarios, such as memory-related tasks. Moreover, while resting-state and oddball paradigms yield comparable results across studies due to their standardised protocols, more complex tasks pose greater challenges for comparison. To address this, it would be beneficial for researchers globally to agree on standardised technical implementations for these tasks, facilitating easier comparison of results and downscaling their heterogeneity. Although some efforts have been made in this direction ^14^, significantly more work is required to achieve widespread standardisation.
3. Expanding on the previous recommendation, we suggest using neurophysiological recordings of cognitive tasks that are minimally obstructed by educational and cultural barriers. While auditory and visual oddball paradigms are an excellent choice for their feasibility and universality ^14^, they do not provide a direct measure of conscious cognitive functions that are typically affected in healthy aging, MCI, and dementia. Among various potential options, music has recently become a viable solution for studying how the brain makes predictions within the context of long-term memory encoding and recognition ^111–113^. For example, in a recent study on aging ^114^, our group demonstrated the functional reorganisation of the brain in healthy older adults while recognizing previously learned music. This task had minimal cultural bias and was straightforward to administer. Despite its simplicity, it provided novel insights into how the aging brain adapts to make accurate predictions, revealing neural differences between younger and older healthy adults even in the absence of changes in behavioural performance. This suggests the significant potential of musical memory tasks for studying aging and identifying novel biomarkers for memory decline and dementia. Future studies should continue exploring this approach and seek other tasks, similar to music ^111^, that are easily comparable across educational levels and cultures.
4. Connected to the previous recommendations, we propose that researchers share EEG, MEG, and M/EEG databases whenever feasible, particularly those using standardised resting-state eyes-closed and oddball paradigm protocols. This practice will enhance the reproducibility of results, facilitate comparison of different methodologies, and reduce the need for new data collections, thereby lowering costs. This is especially crucial today, as current machine learning algorithms can detect fine-grained features in neurophysiological signals that distinguish different groups of participants. Notably, some efforts have already been made in this direction (e.g., UK Biobank ^115^). To emphasise the potential impact, we suggest that significant breakthroughs might already be within reach but have not been realised due to the lack of integrated datasets. Future directions should aim to support data sharing and the consolidation of datasets, while working to make current regulations smoother for easier data sharing and ensuring adherence to ethical and legal standards. This will enable active collaboration and more effective utilisation of existing data, along with bridging the gap between studies from high-, middle-, and low-income countries.
5. It is advisable to conduct more studies on various types of dementia, beyond ADD, due to the underrepresentation of less common forms. Along this line, a further general recommendation would be to enhance the rationale and hypotheses linking dementia types to changes in resting-state and oddball paradigms, as well as other cognitive tasks. Strengthening theoretical foundations and hypotheses could notably improve the field’s quality and the integration with clinical practice.
6. While the search for novel neurophysiological markers is promising, the extraction of numerous EEG and MEG metrics from data has previously posed challenges in comprehensively elucidating the neurophysiology of aging. Addressing this limitation requires a methodological and analytical approach aimed at developing a unified theory of neurophysiology of aging, particularly in standardised procedures. Specifically, while many resting-state studies have examined power spectral density and overall connectivity patterns, a cohesive understanding of systematic changes in brain networks across different frequency bands with aging remains unclear and warrants further investigation. Furthermore, in the context of mismatch negativity responses in oddball paradigms, it would be advantageous to identify the key networks that primarily capture the variance in the multivariate, high-dimensional M/EEG data and to explore how these networks are modulated by aging, thereby enhancing the refinement of existing studies.
7. Most articles reviewed aimed to identify biomarkers relevant for clinical applications in MCI and dementia. However, their primary focus was validating discrimination tools at a group level, which aligns with the standards of scientific publications. While this approach yielded valuable results, its utility for individual clinical use is limited. Indeed, the reliance on biomarkers that necessitate data from entire group populations raises concerns about their value in assessing individual health status. Therefore, while future studies are encouraged to continue drawing inferences at the group level, they should also refine paradigms and analytical methods to enable the use of biomarkers at the individual level. This approach, especially if applied in longitudinal studies, would facilitate their integration into the development of tools suitable for clinical practice.

To sum up, widespread interest in the neurophysiology of aging has quickly grown over time, giving rise to an impressive body of literature on the neural mechanisms of healthy aging, MCI, and dementia. Based on the extracted features, we can conclude that the classic neurophysiology experiment on healthy and pathological aging employs a cross-sectional design, measures spectral features of EEG during resting awake with eyes closed, and comprises at least one population group with ADD. With the predicted increase in global prevalence of dementia, advances in the field of neurophysiology lay the ground for novel diagnostic tools that could be adopted in clinical practice. The implementation of longitudinal designs, standardisation of methodologies, and correlation with established biomarkers represent the key challenges for drastically improving the field.

## Data availability

The code used for computing descriptive statistics and plotting results is freely available on GitHub: https://github.com/gemmaferu/systematic-review-neurophysiology-aging.git.

Data from supplementary tables will be made available upon reasonable request.

## Acknowledgements

The Centre for Music in the Brain (MIB) is funded by the Danish National Research Foundation (project number DNRF117). L.B. is supported by Lundbeck Foundation (Talent Prize 2022), Carlsberg Foundation (CF20-0239), Centre for Music in the Brain, Linacre College of the University of Oxford, Society for Education and Music Psychology (SEMPRE’s 50th Anniversary Awards Scheme), and Nordic Mensa Fund. M.L.K. is supported by Centre for Music in the Brain and Centre for Eudaimonia and Human Flourishing, which is funded by the Pettit and Carlsberg Foundations. We thank Antonio Malvaso for his early contributions to this work.

## Author contributions

L.B., G.F.R. and M.L.K. designed the study. L.B., G.F.R. and M.L.K. planned and implemented the search, screened studies, and extracted relevant data from studies. L.B. and G.F.R. conducted the data analysis and visualisation. L.B. and G.F.R. wrote the first draft of the manuscript. M.L.K. and P.V. provided essential help to interpret and frame the results. All the authors contributed to and approved the final version of the manuscript.

